# A pilot study to define and identify future priorities into *Allocasuarina robusta* recovery as part of a community program

**DOI:** 10.1101/2021.04.19.440386

**Authors:** Matthew W Pearson

## Abstract

The *Allocasuarina robusta* pilot study investigated the process involved to facilitate seed recruitment as part of a threatened species project. Several experiments occurred, each examining a specific attribute in the seed recruitment process. *A. robusta* is a threatened species of national and local significance. The research design would help land managers and communities to conserve *A. robusta*. The investigation aimed to improve seed recruitment in *A. robusta* occurring under natural conditions. The experiment results highlighted several experimental design flaws and identified opportunities to increase community participation as part of the recovery program.

## Introduction

Community involvement is a fundamental aspect for aiding environmental restoration; communities contribute by providing labour and time to assist with natural regeneration or ecological reconstruction activities. McGregor and McGregor (2020) describes the value and the need for environmental restoration and the intrinsic value to the overall community. Communities can drive a project, but communities that manage areas containing a threatened or rare species cannot perform their role without technical information (Gollan et al., 2012; Roger et al., 2020). These communities can direct research through citizen science programs or other community activities by providing detailed local knowledge (Gollan et al., 2012; Roger et al., 2020).

The purpose of revegetation goes beyond the need to augment natural associations with tube stock or direct seeding to protect biodiversity (Breed et al., 2012; Navarro-Cano et al., 2019; Pearson, 2019). Revegetation is to sustain a population by increasing genetic diversity or stimulating the natural recruitment process without the threat from introduced species (Breed et al., 2012). Revegetation or reconstruction techniques have proven helpful for protecting natural communities against further degradation (Breed et al., 2012). While regeneration/reconstruction measures will aid and facilitate environmental protection via a big-picture perspective (Hobbs, 2017). The *A. robusta* recovery program has highlighted the need to establish the seed production areas (Quarmby, 2011). Seed production areas or seed orchards can contribute to biodiversity conservation (Zinnen et al., 2021). The challenge for a recovery program that uses tube stock to ensure species diversity is having the seed of suitable quality and quantity available for use in growing the tube stock (Zinnen et al., 2021). Current regeneration/reconstruction could all be undone by not understanding what occurs at an individual species level (Breed et al., 2012; Pearson, 2019).

Identifying what environmental cues to measure for seed recruitment in *A. robusta* means understanding and testing the hypothesis and significance factors (Janvry et al., 2016; Schmid et al., 2017). Plaisance et al. (2021) described that hypothesis testing and significance testing relate to each other. The relationship can provide misinformation or incorrect inferences when these are not analysed (Robinson, 2018; Schmid et al., 2017). A cornerstone of science is the generation of questions; this may involve creating or reaffirming known knowledge or applying an experimental design differently, none of which will lead to misinformation. Misinformation from experimental design occurs at the execution stage, affecting the results and data analysis (Newman, 2008; Symes et al., 2015).

The reporting of experimental design errors or the generalisation of results should not occur in the opinion of (Robinson, 2018). Undertaking trials is vital for developing appropriate scientific inquiry skills, reducing the possibility of reporting misinformation originating from incorrect experimental design (Symes et al., 2015). A new experimental method or application creates a tendency to focus on procedural components of the experimental design rather than the unexpected outcomes (Hoban & Strand, 2015; Symes et al., 2015). Designing a pilot study should still contain rigour that can test the original research question. As Schmid et al. (2017) explain, the concept of an experimental design identifies and engages with the theoretical aspect of testing.

The theoretical aspects of experimental design and its meaning should remain in context (Newman, 2008). Maintaining the context and reporting of the results could occur by reporting the experimental design limits (Chen, 2010; Newman, 2008). Robinson (2018) concluded that even if negative results, occur these still require reporting to avoid incorrect inferences from single and isolated tests. A fundamental aspect of understanding restoration ecology is knowing what observations are required and why (Pennock, 2004; Robinson, 2018; Schmid et al., 2017). The current investigation occurs in a simulated environment instead of a field study where Schmid et al. (2017) used a homogeneous environment to exert control in the experimental design. Reporting the experimental design needs contextualisation towards the outcome’s size (Hoban & Strand, 2015; Janvry et al., 2016). A pilot studies experimental design involves a degree of scalability where procedures and the testing rigour can be measured (Hoban & Strand, 2015; Schmid et al., 2017). For example, the results from Navarro-Cano et al. (2019) field plot investigation involved changing the scale to maintain genetic diversity in a species. Schmid et al. (2017) extend the concept of scale to measure the time taken for recording the manipulative experimental data and the experiment’s duration.

The pilot investigation aims to demonstrate the importance of experimental design in the restoration ecology. The pilot investigation uses the species *A. robusta*, a threatened species, to test what environmental cues would simulate the populations to regenerate following the parent plant’s death.

## Method

### Study Species

The climatic conditions favouring *A. robusta* is the Fleurieu Peninsula’s wettest parts (Department for Environment and Heritage, 2007). The Fleurieu Peninsula has a temperate climate with moderately wet winters and hot, dry summers (Armstrong et al., 2003). Rainfall in the Fleurieu Peninsula can range from 400 to 1000 mm depending on altitude and aspect (Armstrong et al., 2003). *A. robusta* grows on soils described by the Department for Environment and Heritage (2007) as infertile acidic soils associated with peat bogs. The soil types range from mottled yellow ironstone soils, gravelly duplex soils and sandy glacial outwash soils (Department for Environment and Heritage, 2007).

*A. robusta* occurs in the Kanmantoo bioregion, including the southern Mount Lofty Ranges, Fleurieu Peninsula, and Kangaroo Island (Department for Environment and Heritage, 2007). *Eucalyptus* open forests and woodlands contain *A. robusta* located on the peripheries of wetlands where the mesophytes and hydrophytes meet. The critically endangered Fleurieu Peninsula wetlands communities have legislative protection from the Commonwealth and South Australian State governments (Department for Environment and Heritage, 2007).

*A. robusta*, a threatened species of the Mount Lofty Ranges, is described as a monoecious shrub with smooth bark (Wilson & Johnson, 1989). The branchlets and the scale leaves of *A. robusta* are glabrous, with the immature scale leaves overlapping (Wilson & Johnson, 1989). The female inflorescences produce aggregate fruit from a 3mm long peduncle; these may be sparsely pubescent or sessile on the peduncle (Wilson & Johnson, 1989). *A. robusta* seed description is a samara with seed ranging from 5.5 to 6.0 mm in size (Wilson & Johnson, 1989).

Pollination in Casuarinaceae occurs by the wind; the bracteoles develop into a fruit that contains a single winged samara seed (Wilson & Johnson, 1989). The female inflorescence develops a woody cylindrical infructescence consisting of whorls of tightly appressed hairs of enlarged floral bracteoles (Wilson & Johnson, 1989). Growth of the floral bracteoles becomes part of the formation of aggregate fruit in *Allocasuarina*. The aggregate fruit is initially hairy and then develops two woody valves with the seed filling the cavity (Wilson & Johnson, 1989). *A. robusta* stores the seed above ground and then releases seed through an environmental trigger (Quarmby, 2011).

### Seed Collection

*A. robusta* seed collection occurred from 11 sites from the species distribution in the southern Mount Lofty Ranges in South Australia from Hindmarsh Tiers (Figure 1). The seed collection occurred from September 2017 to February 2018. The collection technique involved collecting aggregate fruit by hand from branches near the base close to the main stem and branches that had not hardened but were still flexible. These sites had additional tube stock planted from other populations within the *A. robusta* range. Sites selection occurred along an east to the west gradient, and two outlying populations on the north and south. Two seed collections occurred; the first a mixture of all 11 sites to form a composite seed collection to test the different simulation techniques. The second seed collection was seed collected to investigate the populations at an individual level.

**Figure 1:**
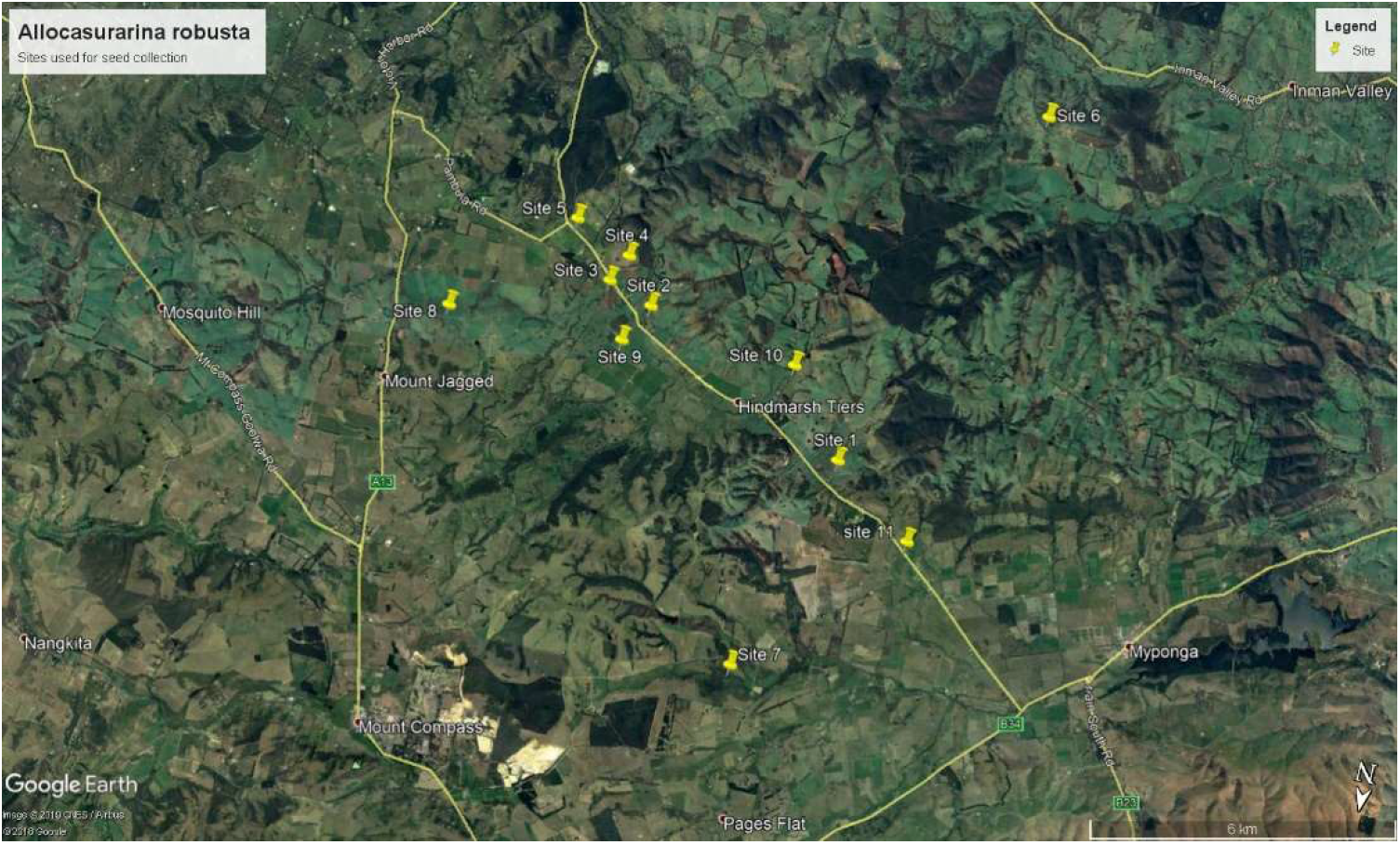
Seed collection sites of *A. robusta*

### Seed examination

Different age *A. robusta* seed was collected and examined. Aggregate fruit from different positions on the *A. robusta* represents a cross-section of age to avoid environmental variability. In each population, sampling occurred on 10% of the population. The 10% sampling relates to the conditions stipulated in the permit from the Department of Environment and Water in South Australia. Each population had a variable number of *A. robusta* from 10 to 112 individuals.

Examination of seed from various populations was under a dissecting microscope. The initial visual assessment enabled the development of a seed characteristic list. Several characteristics initially included were removed due to being reflective of seasonal variation in the seed. Cochrane et al. (2015) defined that seasonal variations are a cause that affects the seed size (width and length) and the seed’s plumpness.

The names assigned to the *A. robusta* metapopulations are road names or nearby local features, i.e., Hindmarsh Tiers Road had two metapopulations with one sample allocated the road name Hindmarsh Tiers and the second sample named after a nearby local feature Hindmarsh Falls.

The *A. robusta* characteristics list used.

#1. The geometry of the seed and samara is/

1. seed and the samara are symmetric/
2. seed and the samara are asymmetric/
#2. Midrib in the samara/

1. shows no signs of tapering/
2. shows tapering away from the seed/
#3. Seed surface characterised by being/

1. entire with surface pitting/
2. entire without surface pitting/
#4. Samara colour is/

1. brown/
2. clear/
#5. Seed colour is/

1. brown/
2. black/
#6. The colour pattern between the samara and seed is/

1. ragged without any fading towards the end of the samara/
2. entire and fades towards the end of the samara/
#7. The surface texture of the seed is/

1. rough/
2. smooth/
#8. The placenta connection between seed and fruit/

1. is retuse with smooth edges/
2. is entire with smooth edges/
#9. The shape of the seed is/

1. generally square/
2. generally oval to round/
#10. The shape of the end of the seed is/

1. obtuse to acute in shape/
2. rounded to ovate is shape/
#11. The seed has a colour marking which gives it an appearance of/

1. a single colour without striations (stripes)/
2. striations (stripes)/
#12. Number of teeth/

1. 9 or more/
2. 8 or less/
#13. Number of protuberances on the Aggregate fruit are/

1. single on the back of the bracteole/
2. several on the back of the bracteole/
#14. The shape of branchlets are/

1. rounded or subangular/
2. angular/
#15. The phyllichinia has/

1. a shape that is rounded or subangular/
2. a central groove/
#16. Aggregate fruit are on/

1. pedicels less than 2 mm long or sessile/
2. peduncles 3 – 12 mm long/
#17. Length of teeth/

1. 0.6 – 1.5 mm long/
2. 0.3 – 0.5 mm long/
#18. Teeth bases are/

1. not overlapping/
2. overlapping/
#19. Bracteoles of fruiting cone/

1. thick and convex, often with separate angular or divided protuberances/
2. relatively thin and without any dorsal protuberances/

### Competition / Nursery

To determine if the presence of either the Burr medic (*Medicago polymorpha*) and Cocksfoot (*Dactylis glomerata*) contributed towards acting as a surrogate nursery to allow for the establishment of *A. robusta*, a total of 30 punnets sown with the introduced species, i.e., 30 punnets of *D. glomerata* and 30 punnets of the *M. polymorpha*. Fifteen were randomly selected and sown with *A. robusta*, which gave a combination of 15 *A. robusta* and *M. polymorpha* punnets and 15 *M. polymorpha* only punnets. A repeated method occurred involving replacing the *M. polymorpha* with *D. glomerata*. The experiment design contained fifteen punnets to act as control with *A. robusta* without any form of competition. Punnets remained in growth boxes for 100 days. The sowing of *A. robusta* did not occur until day 20. The 20-day lag time was to allow the *D. glomerata* and *M. polymorpha* to establish. Response variables were the time of germination and survivorship over 100 days.

### Surface litter

Observations in Quarmby (2011) noted that dying *A. robusta* resulted in seed released from the canopy. The experiment designed was to mimic conditions of natural seed recruitment in the absence of fire. The experiment used a square squat pot to ensure variability of litter depth while maintaining a proper soil depth to allow a seedling to establish. Square squats (470ml) from Garden City Plastics (https://www.gardencityplastics.com/) filled with 100ml of growing media and seed were sown at a density of 30 seed per pot to represent a natural seed rain. The first treatment had a layer of seed placed on the surface litter to replicate natural seed rain. The second treatment placed the seed on the interface between the surface litter and the growing media to represent seed released and covered with leaf litter (10 mm of leaf litter). The third treatment buried the seed into the litter to a depth of 25 mm, to represent a historical seed release event. The leaf litter used was from *Eucalyptus cosmophylla* (Cup Gum), with the leaves collected from nearby roadside reserves. These were then washed, dried, and exposed to U.V. light to reduce any possible contamination on the leaves. A fourth treatment sowed the seed on the surface of the growing media without any leaf litter. Each pot was given a number and then assigned to a random location within a four-block experimental design. Each treatment replication occurred eight times, resulting in 24 square squats used with six square squats per block. Response variables were the time of germination and survivorship over 100 days.

### Growing Media

Quarmby (2011) described the *A. robusta* populations’ location on the perched water table, forming the Fleurieu Peninsula swamps. The experiment’s design was to determine which soil type would be conducive for *A. robusta* seed recruitment. The experiment used commercial growing media, which varied in the sand and organic matter ratio. Four treatments varied soil media. These were propagation sand (http://www.brunnings.com.au/propagating-sand-5l.html), orchid mix (https://www.searlesgardening.com.au/products/category/OTNTNCPH-speciality-range/LDEB--searles-dendrobium-orchid-mix-30lt), all-purpose growing media (https://www.debco.com.au/products/all-purpose-potting-mixes/debco-premium-potting-mix) and natural soil was determined using a mixture of the soils collected over three sites. Sites were Stipiturus Conservation Park (Site 7), Mt Billy Conservation Park (Site 5) and Hindmarsh Falls (A water reserve managed by the Yankalilla District Council, Site 10) (Figure 1). Treatment of natural soil occurred to avoid any competition using solarisation and heating at 200°C for 20 minutes to sterilise the soil. Sowing of twenty *A. robusta* seed into each punnet occurred. The experiment conducted was a single block design. Response variables were the time of germination and survivorship over 100 days.

### Heat Intensity

Fresh cones of *A. robusta* were collected, bagged and then exposed to 100°C for five minutes to allow the fruit to release the seed. When the seed was released, the seed was batched into six groups then exposed to 100°C for various durations.

Exposure length was at intervals of 0 minutes, 2 minutes, 4 minutes, 6 minutes, 8 minutes, and 10 minutes. The seed sowing occurred in sand-based growing media. Each of the timed heat exposed groups of seed was replicated four times with thirty seeds sown in a 400 ml punnet. All six treatments were allocated a number and then randomly distributed into a growth box. Response variables included germination over the next 100 days and survivorship.

### Heat Shock / Smoke

To simulate the impact of fire. Seeds sowing occurred in wet growing media where treatment was applied and then placed in a three-block design with each treatment replicated three times. Treatments of the seed in situ of the potting growing media included heat shock, smoke water, a combination of smoke water and heat shock and a control. The process of simulating heat shock to the seeds occurred by applying boiling water to the seed to act as a form of heat shock (Mackenzie et al., 2016). The experimental design simulates a fire’s effects on the seed described by Mackenzie et al. (2016). Heat shock does not necessarily mean exposure to fire in the form of flames but the heat generated by the fire (Mackenzie et al., 2016; Pounden et al., 2014). As *A. robusta* seed protected in the fruit from any direct fire impacts (Burrows & Middleton, 2016). The smoke water treatment consisted of using a commercial product from Suregro (http://www.suregro.com/product/regen-2000-smoke-master-liquid-5-litre/) and making up a solution of five litres (rate 0.1: 10 ml water) and applied to punnets through overhead watering

### Seed depletion

To measure the seed bank depletion for the *A. robusta*, thirty cones collected from the west to the east gradient at Hindmarch Tiers (Figure 1). Mixing of the cones occurred to create a random source of seed. Grouping fruit occurred by site with six cones per paper bag, providing a comprehensive collection of five sites. Cones were dried and stored with temperatures ranging from 16.7°C to 19.9°C, with relative humidity ranging from 84% to 88%. The first bag of seed was sown two weeks after collection, with each subsequent bag sown every four weeks after the initial sowing. Sowing occurred at a rate of 30 seeds per 400 ml plastic punnet with each treatment replicated four times then placed in one of four blocks in the growth boxes. The growing media used was a commercial growing media that was sand-based, including composted organic matter (https://www.debco.com.au/products/all-purpose-potting-mixes/debco-premium-potting-mix). These were monitored daily for germination and survival over 100 days.

### Population viability

Seed collection occurred at eleven of the twenty-four populations of *A. robusta*. Seed collection occurred from individual *A. robusta* with 10% of a population sampled with ten cones selected from each individual. Air drying of the *A. robusta* cones without any environmental controls. Each of the bags was labelled dated, and site information included. The seed was sown two weeks after collection. The seed was sown in 200 ml punnets using sand-based growing media. Each punnet had thirty seeds sown, with each *A. robusta* collection site replicated four times. Watering of punnets occurred before sowing and on completing sowing of the blocks used in the growth boxes.

### Growing condition of the seed treatments

Seeds were sown in 200 ml commercial nursery punnet using sand-based growing media unless specified elsewhere. The seed germination occurred in a growth box modelled on the progradation bed designed by Sage Horticulture (https://www.sagehort.com.au/propagation-equipment/propagation-beds/PROPAGATIONTRAYHEATMISTENCLOSURE) using PSI lighting with purple globes. The growth boxes used were 149L clear plastic storage boxes with lighting fixed to the lid. Each box was filled to a depth of 20 mm of playground sand with a heating pad (Medium Seahawk Heat Pad) then covered with a further 20 mm of sand.

The application of heat pads if a predicted frost was forecasted. Daily seed data collection occurred, and watering occurring every second day of approximately 150 ml of water applied (via a mist system) to each treatment. Other data observational data included recording the seedling’s survival; assigning a survival category, was cotyledons only or visible central stem development with scale leaves. The seed collected from *A. robusta* along roadside corridors occurred from random plants. The seed collection contained a limitation of no more than 10% of fruit collected from any individual and no more than 10% from a population. To meet the South Australian Scientific Permit (A26769-1) from the Department of Environment and Water requirements. Watering used for the punnets consisted of using Adelaide mains water without any purification or treatments. No additional fertiliser or plant growth regulators were used to establish the *A. robusta* seed unless it was part of the experimental design.

### Statistical Analysis

Descriptive analyses occurred using and XLSTAT (Mélard, 2014). The descriptive statistics included looking at the data’s central tendency (mean median, mode, standard deviation, and variance). Each experiment’s design was a balance block design to enable ANOVA to identify statistical significance; if this occurred, further examination of the data occurred using RStudio (R Development Core Team, 2010) for linear regression and frequency distribution plots. Single variables formed the basis of the investigation as each of the processes impacted germination due to being a pilot study; no combinations of data occurred.

The seed examination used a dissecting microscope at x45 magnification. Data analysis from collected data occurred in Delta Ver1.02 (Dallwitz et al., 2013) and PAUP Ver.4.0a (Swofford, 2001). PAUP Ver.4.0a analysed data as an initial branch and bound analysis tree based on parsimony’s informative characters. Topological constraints and trees that were unrooted were turned off.

## Results

### Seed examination

Examination of seed collected from *A. robusta* occurred before use in the manipulative trials, from the criteria provenance and implications for environmental restoration. The character list design will identify any observable differences in morphology and establish a seed provenance for *A. robusta*. When the parsimony data generated in PAUP Ver.4.0a, the results showed insignificance for consistency index (0.5926) and homoplasy index (0.4783) towards morphology from the sampled populations.

### Competition / Nursery

*A. robusta* sown beneath the *D. glomerata* had only single germination, comparable to the A. robusta seed sown without competition with single germination. Whereas the *M. polymorpha* / *A. robusta* mixture had 12 seed of *A. robusta*, which germinated. The number of seeds germinated in the growth box was not enough for statistical analysis, but they show some interesting relationships with neighbouring vegetation. Not all seed germinated simultaneously, with the first seed germinating on day 14 and the last seed germinating on day 55 (Table 1) in the *M. polymorpha* / *A. robusta* mixture. The results for *D. glomerata* being only single germination occurred on day 23. During the 100 days, only one seedling died.

**Table 1:** Germination of *A. robusta* with *M. polymorpha*

| Day | Germination | The germinated seedling withered at the cotyledon stage |
| --- | --- | --- |
| 14 | 4 | 0 |
| 14 | 3 | 0 |
| 27 | 0 | 1 |
| 27 | 2 | 0 |
| 35 | 2 | 0 |
| 55 | 1 | 0 |

In Table 1, *A. robusta* competition with the *M. polymorpha* only occurred in two of the replications, with the remainder having no *A. robusta* germination. Observation on these treatments, the punnets with *M. polymorpha* appeared to be wetter compared to those with *D. glomerata*. Another observation between the two introduced species is that the *M. polymorpha* also had a less aggressive root system than the *D. glomerata*. From Table 1, germination occurred over 41 days, with germinations decreasing as time progressed.

### Surface litter

Eleven seeds germinated over 100 days. Germination began on day 36, and by day 88, it had concluded. Seed buried to a depth of 25mm recorded no germination, yet the seed sown at the interface between the surface litter and the growing media began to germinate on day 65 and concluded by the 88^th^ day. Seed sowing occurred to simulate seed rain with germination begin on day 36 and had concluded by day 85. Simulated seed rain without leaf litter had a more significant number of germinations than when the seed covered by leaf litter (F-value 2.631, DF 6, St. Dev. 19.292, P=0.061).

### Growing Media

A majority of the germination occurred in the propagation sand, with germination beginning on day 19. Longer germination times occurred on the general-purpose growing media, with limited germination occurring on the composted orchid growing media. In the natural soil mixture, only a single seedling appeared on day 42. The propagating sand had a significantly higher germination rate (F-value 4.494, DF 17, St. Dev. 5.916, P=0.009). The general-purpose growing media (10 germinations F-value 4.494, DF 16, St. Dev. 1.182) and the composted orchid media (three germinations F-value 4.494, DF 18, St. Dev. 1. 182) produced the same results (P=0.01). Even though the composted orchid growing media did have statically comparable results, the number of observed germinates were less.

### Heat Intensity

Exposure of the seed to 100°C at 2 minutes intervals and finishing at 10 minutes resulted in 36 germinations. From the observation notes, 4-minute exposure time produced the highest number of germinations compared to treatments. Germination began on day 25 and concluded by day 80. From the observations, the more significant exposure time to heat resulted in longer germination times, with the adverse occurring for a reduction in heat exposure. The control treatment began germinating after 19 days. The observational results indicate that 4 minutes at 100°C is optimum for maximising germination.

### Heat Shock / Smoke

Only three of the punnets produced any germinations over the 100 days. Seed treated with smoke water began germination 21 days after sowing. From the treatment, only three of the four produced results and one germinant recorded in the control treatment (Table 2)

**Table 2:** Treated *A. robusta* seed germination responses

| Treatment | Number of germinations |
| --- | --- |
| Smoke Water | 3 |
| Smoke Water and Heat Shock | 3 |
| Heat Shock | 0 |
| Control | 1 |

**Table 3:**
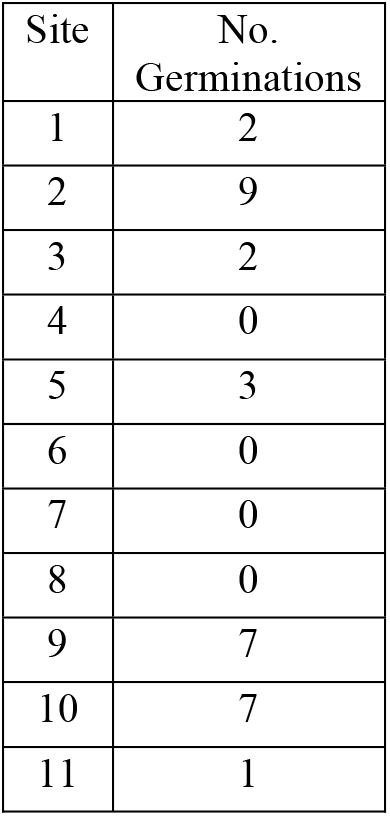
Table results from the Population viability experiment.

| Site | No. Germinations |
| --- | --- |
| 1 | 2 |
| 2 | 9 |
| 3 | 2 |
| 4 | 0 |
| 5 | 3 |
| 6 | 0 |
| 7 | 0 |
| 8 | 0 |
| 9 | 7 |
| 10 | 7 |
| 11 | 1 |

From Table 2, the two-treatment exposed to smoke water had much higher germination compared to heat alone or no treatment.

### Seed depletion

Sowing seed and storing the seed at an average ambient room temperature of 23°C produced only one germinate after 30 days. No other germinations occurred for the next 180 days. The seed was sown every thirty days for up to 120 days, where sowing ceased, and observation continued to day 180.

### Population viability

Examining eleven populations from west to east of the *A. robusta* over 100-day germination, only 31 germinates occurred. Population 2 produced 29.03% of germinants. Two other population produced seven germinants (22.58%), and other populations produced between 0-3 germinates. From a geographical interpretation of the data, all three sites with a high percentage occurred in the same valley system. Those sites which recorded no germination were in smaller and isolated valley systems.

## Discussion

From the investigation conducted, it would be easy to conclude that the experimental design was flawed. However, the design flaws provide several vital points for directing future research and identifying the parameters requiring alteration (Chen, 2010; Evans et al., 2020; Newman, 2008). The notion of reporting results that are flawless or consistent with theory or expectation would weaken application to a meta-analysis (Plaisance et al., 2021; Winkle-Wagner & McCoy, 2016). Roux et al. (2017) and Winkle-Wagner and McCoy (2016) identify how scientific inquiry does not always occur in ideal circumstances but managed through appropriate variable controls. Reporting on which controls to manage or which variables should be analysed has, as Winkle-Wagner and McCoy (2016) indicated, resulted in selective reporting. When considered holistically, the scientific inquiry process and the reporting of either negative or positive results is necessary (Plaisance et al., 2021; Roux et al., 2017). When a particular outcome does not occur in the investigative process, these areas allow learning to occur from reflecting on the experiment’s hypothesis and design (Chen, 2010; Evans et al., 2020; Symes et al., 2015). The current experimental design contained elements from Barritt and Facelli (2001), Abihudi et al. (2020) and Huang et al. (2021). The basis for scientific inquiry is the generation of new information and then explaining the results in the context of the current theory (Plaisance et al., 2021; Roux et al., 2017). By piloting several methods can help identify and redefining the research question, which was the case for the current investigation. The aspect of piloting an experimental methodology and then reporting on the outcome is not new, as Evans et al. (2020) used to refine and identify research areas on seed recruitment. The results’ perception indicates a general lack of significance from the current investigation, which supports the need to ensure pilots experiments or trials occur before larger experimentation is conducted.

From the simulated leaf litter effect on *A. robusta*, the results lacked significance (P=0.061). However, from a similar type of investigation performed by Barritt and Facelli (2001), the leaf litter would impact seed germination. Barritt and Facelli (2001) discussed how simulated, or natural forms would not hinder seedings’ emergency. The *Casurarina* litter is like *A. robusta* litter would be loose and provide no physical effect on seedling emergency (Barritt & Facelli, 2001). In the litter experiment conducted for the *A. robusta* and Pandey et al. (2020) investigation, both experiments only measure a single factor being a seedling emergency. Pandey et al. (2020) indicate that further investigation required on the impact of the seedling emergency, including light availability, competition, and soil community (e.g., fungi). From the experiment conducted, the p-value produced was not less than 0.05 and in Schmid et al. (2017) review, such a result would require retesting or undertaking the experiment again. Nerveless in Schmid et al. (2017) context, the results do not provide confidence towards the significance or the effect size. The lack of confidence from the results produced could, as Robinson (2018) discuss, arise from a lack of replication, but the results contain several similarities to Barritt and Facelli (2001) investigation.

Replication can improve experimental precision, but replication does not always solve experimental design issues (Schmid et al., 2017). The collection of samples to examine seed morphology of *A. robusta* occurred on an east to west transect through the population. Data collection on seed morphology may not be an accurate ecological indicator of the species’ health. Additional data could be collected, including other morphology features, including phenological data related to the species. The addition of phenological data would explain how or when *A. robusta* seed development begins (McDonough MacKenzie et al., 2020). Phenological data would not resolve the sampling aspect, but McDonough MacKenzie et al. (2020) discuss how phenological data supports taxonomy and seed provenance questions. The caveat that needs to be applied is the number of sampling points, and the number of aggregate fruits selected could allow pseudoreplication to occur (Schmid et al., 2017). Phenological data is not solely focused on when a species flowers but can include when a species is actively growing. Phenological data can increase the taxonomic breath in an investigation by providing supportive information for taxonomy (McDonough MacKenzie et al., 2020). Analysing *A. robusta* seed morphology contained insignificance from the parsimony analysis. McDonough MacKenzie et al. (2020) indicates that phenological data can resolve the parsimony analysis’s insignificance. The application of phenological data would increase the diversity in sample data. Applying phenological data would increase data diversity, but care needs to be applied equally to ensure a suitable and representative sample size occurs.

Pennock (2004) indicates that when experimental design lacks sample diversity, it can give rise to pseudoreplication. The investigation focused only on one species (*A. robusta*), but the seed morphology experiment included comparing *A. robusta* relatives seed. Chen (2010) explains how this allows researchers to examine and develop an alternative research question. *A. robusta* surface litter results were comparable to Barritt and Facelli (2001) investigation based on the current investigation results. Barritt and Facelli (2001) used a different species along with a different environment habitat. However, future investigations should compare the threatened *A. robusta* to a species of least concern from the same environment. The same type of concept occurred in Abihudi et al. (2020). Identifying which species to use for the species of least concern could come from Pearson (2020) investigation. The experimentation process tested heat shock, smoke, and seed age can provide vital information for managing a threatened or rare species. Considering Abihudi et al. (2020) investigation, the data collected in this investigation, coupled with a common species, could indicate a species’ conservation trajectory in the Fleurieu Swamps. Germination of *A. robusta* was low in the current investigation. However, Dwyer (2017) conducted a study using an *Acacia* species found survival of seedling were low as 0.9% this could represent hundreds of thousands of seeding facilitating the species survival on a per hectare basis. In this investigation a single germinant occurred, meaning it would be difficult to come to the same conclusion as Dwyer (2017) on the number of seedlings required to produce a sustainable population.

Comparing a threatened species and a common species occurred in Abihudi et al. (2020). Huang et al. (2021) extended the comparative concept to include an introduced species. The current investigation investigated the competition/nursery effect, which produced single germination with *M. polymorpha*. The assumption was that the competition for resources would only occur through the interaction between a native species and introduced species. Huang et al. (2021) indicate that competition could occur between two native species. Like Huang et al. (2021), the competition/nursery effects occur through the time taken for germination and the speed at which the species establishes. Observations from the current investigation can be related to concepts discussed in Catterall (2019). Catterall (2019) discusses and the role of nurse plants in restoration ecology; the current results will require further investigation to be more conclusive and definitive. Barritt and Facelli (2001) and Pandey et al. (2020) several related factors that can affect germination (i.e. the interrelationships between a nurse plant and the study species), all of which require further investigation. Understanding the role or relationship of a nurse plant between *A. robusta* will require further investigation. Lozano et al. (2020) explored the nurse plant relationship and role in the recruitment process. Lozano et al. (2020) identified that the main contributing factor for establishing a nurse plant relationship is soil, but in *A. robusta*, the evidence collected is inconclusive and requires further investigation. The current investigation had single germination, which poses several questions, did this occur by chance or was it through soil amelioration, as was the case in Lozano et al. (2020). Alternatively, a single germinant’s survival reflects *A. robusta* long-term recruitment strategy, which occurs in other species (Dwyer, 2017; Navarro-Cano et al., 2019). Navarro-Cano et al. (2019) investigated occurred in a different genus by examining the species’ long term recruitment trajectory.

The reporting of the investigation results and the comparison with other investigation (i.e. Abihudi et al. (2020) and Navarro-Cano et al. (2019)) requires a caveat to avoid misinformation to give it a theoretical basis. Palmer (2000) termed investigation, which used different systems and species quasi replication. Identifying and comparing data against other species or different systems is often used, or comparative analysis from a data subset used as a means of justification (Palmer, 2000). Palmer (2000) would not entirely dismiss the role that quasi replication can have in science. Chen (2010) identified that comparing but not analysing the investigation is not a true reflection of the hypothesis. For instance, the design of the heat intensity and heat shock/smoke experiments should demonstrate a significant germination event post fire for seed recruitment in *A. robusta*. The reported results were not conclusive to support or dismiss the hypothesis. A simulated fire used in the experimental design contained no comparative analysis with another species. Comparability between the current investigation and the previous experiment (Pearson, 2021) could not occur due to the differences in the methods used and design used. Cury et al. (2020) and Burrows and Middleton (2016) compared species under the same condition, which provided meaning and a theoretical context. In the current investigation, the absence of a comparison between *A. robusta* and another species provides an opportunity for further investigation.

Replication needs to occur under the same conditions, and while comparison between common/threatened or threatened/invasive can occur. The results from the current investigation examined the metapopulations of *A. robusta*. The results were not conclusive; the appearance of pseudoreplication could occur; as Schmid et al. (2017) describe, the hypothesis was to determine the viability of metapopulations for natural regeneration. The method selected was like the method used by Costa e Silva et al. (2019) for examining the provenance of a Tasmanian *Eucalyptus* species. Costa e Silva et al. (2019) compared *Eucalyptus* provenances through replication within a controlled growing environment and a common garden experiment when reflecting on *A. robusta*, which occurred only in a controlled growing environment. The current investigation requires a comparison to aid the recovery of *A. robusta* further.

From the data collected, several opportunities exist which can further aid the recovery of *A. robusta*. The community’s role in collecting data or identifying new metapopulations of *A. robusta*, particularly on private land identified by Quarmby (2011), should occur. A strategy described by Breed et al. (2012) would provide the ideal means to achieve success through a tube stock program to improve seed recruitment practices. The caveat placed on the strategy needs to be evidence-based for the ecological community and each species used (Breed et al., 2012). The use of seed orchards to restore can have an environmental cascade effect in a community or ecotype from the management practices or the ecological effects of the material used Zinnen et al. (2021). From the investigation conducted, the results were inconclusive or did not address the role of seed orchards. Zinnen et al. (2021) review identified several risks from using seed orchards, including the loss of genetic diversity. The loss of genetic diversity can mean that a population would become more homogeneous over time and may not contain the resilience required for the effects of climate change Zinnen et al. (2021). The investigation results, while inconclusive on the effects of temperature on the exposure to temperature. The loss of a population or the re-establishment of a population following a controlled disturbance event contains various implications for a community-driven project. A seed orchard, as described by Zinnen et al. (2021), would resolve some issues. These events can act as an enabler for allowing introduced species to invade or allow the colonisation of more aggressive endemic species (Davies & Boyd, 2021).

The role of introduced species would have in acting as a nursery plant occurred as part of the pilot investigation. The result indicates a degree of support towards the case studies from Catterall (2019). A different viewpoint provided by Broadhurst et al. (2017) is that the selection of propagules from a seed orchard may reflect the establishment of the species. An aspect which the investigation with further data collection may indicate provenances in *A. robusta* could occur. If provenances have formed, these may contain traits that could facilitate the re-establishment of the species (Davies & Boyd, 2021). Taking the provenances concept can address the concerns of genetic diversity, as Zinnen et al. (2021) identified. Selecting advantageous material of desired species means the quicker establishment reduces the impact of introduced species (Davies & Boyd, 2021).

The establishment of the *A. robusta* and its relationship with a nursery plant, while explored in the current investigation, did not produce statistically significant results. While had some support towards Catterall (2019), the incidental observation made of the differences in root architecture could impact germination and survival of the *A. robusta* seedling. Strumia et al. (2020) considered root morphology and how this would translate to seedling establishment occurred. As *A. robusta* grows on the wetland environment’s peripheries, the hydrological changes are similar to the investigation (Strumia et al., 2020). Understanding how groundwater quality can affect seeding may be imperative for seedling survival. The results from Strumia et al. (2020) can extend to include other soil properties that may impact seedling survival, e.g. structural degradation. Davies and Boyd (2021) identified speed of establishment would be one character for establishing provenances, but soil’s relationship could be another.

The selection of seed material contributes to the current management process undertaken of reconstruction with tube stock. Other management practices identified in Quarmby (2011) implemented, including a controlled ecological fire, occurred. The effect of the fire as a means to simulate seed recruitment has occurred in previous investigations (Bradshaw et al., 2018; Pearson, 2021). The results of Pearson (2021) did indicate fire may not contribute to population recruitment. The linkage between the two requires further examination by considering other attributes of the seed. An investigation conducted by Soares et al. (2021) points to several considerations that could apply to *A. robusta* through measuring the seed water content, fire intensity and seed store proteins when exposed to fire. The aspect of fire intensity and the regularity of fire events can impact the community that develops. From the investigation, the consideration of specific fire cues occurred. Seed exposure to heat or comparison between heat and smoke products form part of the investigation. As the results were inconclusive, other attributes need considering. For instance, Nation et al. (2021) investigation demonstrates an extension of the current investigation concept. Nation et al. (2021) combines fire and the placement of the fruit in the surface fuel. The concept applied by Nation et al. (2021) equally applies to *A. robusta* because the fruit can open through damage to the branch or the parent plant’s death.

Nation et al. (2021) concluded that surface fuels need to contain a mixture of fuel types to protect and stimulate germination. The relationship of heat through undertaking a heat shock investigation showed no conclusive results. From the investigation conducted by Ibañez Moro et al. (2021), exposure temperature was much higher. What differentiates the current investigation and Ibañez Moro et al. (2021) exposes the seed directly to heat shock. Understanding the relationship between heat shock, dormancy and seed stored above ground has had little investigation. Investigations like Ibañez Moro et al. (2021) and Daibes et al. (2021) focus on heat shock related to seed stored in the soil seed bank. The investigation can contribute to the further understanding of seed storage in A. robusta through the experimental design used. The data collection methods used by Daibes et al. (2021) can apply to *A. robusta*. Ibañez Moro et al. (2021) and Daibes et al. (2021) experimental design reinforces the need to ensure a comparison between species can occur. The aspect of compassion between species related to or other serotinous species occurred in Ibañez Moro et al. (2021) and Daibes et al. (2021); if selecting other serotinous species, the experimental designed similar to Soares et al. (2021) can enable comparison between serotinous species can occur.

Soares et al. (2021) examined a serotinous species and concluded that while the species can adapt to climate change, this may not mean the species will maintain the same seed viability level under climate change. For *A. robusta*, further examination of the seed viability and the biogeography beyond what undertaken by Pearson (2020) required to include rainfall and other associated climatic data. Trezise et al. (2020) recent investigation would contradict the findings of Soares et al. (2021); even so, the results from Trezise et al. (2020) offer a different perspective to *A. robusta*. From Trezise et al. (2020), the results would indicate the type of swamp inhabited by *A. robusta* would be drier than those studied by Trezise et al. (2020). Presently knowing if the swamps are different require extending the biogeographical study undertaken by Pearson (2020). In Trezise et al. (2020) and McCaw and Middleton (2015), while no *A. robusta* recorded, there is some evidence to suggest fire would facilitate seed recruitment, whereas, in the current investigation, this would be inconclusive due to the poor germination to test statistically.

The factors contributing to *A. robusta* poor germination did not occur as part of the investigation. In hindsight, this may provide greater depth to explain why germination was significantly lower when environmental treatments were applied. In the investigation, a classification system to aid in identifying seed provenances occurred. The further development of the provenance classification based on seed morphology would help provide insight into the seed traits and seed viability. The investigation undertaken by Guerrero et al. (2021) environmental factor had a significant role in impacting the seed viability. The role of the seed coat provides a protective barrier to protect the future embryo (Guerrero et al., 2021). However, the quality of the seed coat, as Guerrero et al. (2021) indicates, is affected by the environment, which can reduce the seed’s germinability and viability. In the current investigation, exploration of differences in seed consisted of studying discrete populations identified by Quarmby (2011).

The concept of seed orchards, which Zinnen et al. (2021) identified, would facilitate the restoration of species or communities. Two issues raised by Zinnen et al. (2021) was quality and quantity of seed. Having the seed available to augment natural regeneration would require storing seed in an off-site seed storage facility. These off-site seed storage facilities can affect the seed’s viability, which can relate to the storage environment (Uddin et al., 2021). The environmental factors for seed storage facilities would require further investigation. Depending on the temperature, seed moisture content, and storage container, the storage environment could impact the seeds’ germinability (Uddin et al., 2021). Quarmby (2011) identified that restoration work with tube stock would be advantageous for the recovery program. Quarmby (2011), as part of the translation, plan justification, identified that *A. robusta* readily germinates in a nursery environment. The current investigation did not occur when undertaken in simulated field conditions or from the investigation (Pearson, 2021). Storage conditions of condition, mainly how the loss of water from the seed, can play a critical relationship to species’ germinability (Uddin et al., 2021). Success and seeing success when involving the community can be vital to ensure the recovery program success.

The current management of *A. robusta* involves intervention using fire in an ecological burn or planting tube stock. The community-driven restoration project would facilitate the recovery of the *A. robusta* (Pearson, 2019; Quarmby, 2011). However, our knowledge of the species phenology regarding aggregate fruit formation is incomplete (Quarmby, 2011). From the current investigation, various assumptions occurred regarding the simulated fire to mirror those of a spring ecological burn that may be detrimental to the species health. Identifying a suitable timing for reconstruction work was critical not only for the species survival but for associated work, including weed control (Yamada et al., 2021). When competition is minimal, tube stock planting will facilitate the establishment of *A. robusta* identified in similar investigations (Yamada et al., 2021).

From the current investigation, while no insight or direction to recover *A. robusta* provided, some crucial advice for practitioners in the field of restoration ecology given. Volunteers/communities are motivated for a range of reasons to participate in environmental programs; successful programs require to have beneficial outcomes (Hobbs, 2017). The two key areas from the results related to fire and seed collection demonstrate fickled success. Science might identify success through statistical significance, but the measure of a successful program may be different for the community (Hobbs, 2017). The current investigation had a rather noble goal to identify success in the *A. robusta* recovery project of a self-sustaining population. The pilot trials conducted had inconclusive results. Examining the results and the concept of how a community might interpret the results demonstrate the importance as Hobbs (2017) discusses that restoration is n ongoing process that needs to adapt and learn from the actions that have occurred.

How a community perceive the investigation can translate into the level of participation and involvement which might occur going into the future. Communities can have a vital role in the success of a recovery program or assessing an ecological community’s quality, mainly when incomplete knowledge is an issue (Leduc et al., 2021). Gaining the knowledge can occur through a variety of process, the use of primary knowledge, (e.g. a landholder) or the use of the community to collect data support scientific endeavours (Gollan et al., 2012; Leduc et al., 2021). The relationship between community-driven action and science can support the recovery of *A. robusta*. From the investigation conducted, further investigation into seed recruitment of *A. robusta* is still required. Breed et al. (2012) discuss genetic diversity and variability as part of the seed collection strategy. Through population genetics, this would aid in understanding if areas of seed provenance exist or not. The genetic data can aid and inform the community on how to progress the recovery efforts. Informing the professional’s recovery efforts is not a one-way process, but a twoway process as the community can inform the professional to direct and support the research (Gollan et al., 2012). Gollan et al. (2012) identified that data collected from volunteers (community) is comparable to professionals’ data. Using the findings from Gollan et al. (2012) investigation and applying them to the *A. robusta* recovery project could mean that replication for common garden experiments could occur in various field locations where citizen science monitoring can occur. For instance, the investigation involved a competition/nurse plant or leaf litter could be an ideal project that the community could undertake as a common garden experiment in collaboration with the professional. The collaborative work undertaken as part of a citizen science project as part of the community engagement can produce valuable data aiding the recovery efforts of a threatened species (Roger et al., 2020).

Defining the role of citizen science is essential for ensuring the success of the program. Community involvement in threatened species management would involve ensuring that some degree of quality control would be occurring over the data collection, which can occur through benchmarking activities for data collection (Gollan et al., 2012). Design consideration for increasing community involvement would, as Gollan et al. (2012) discuss, need to have simple and easy to use data collection tools. Gollan et al. (2012) described several case studies where presence/absence data collection tools or flipbooks to identify species present as the most effective tools. Gollan et al. (2012) ideas can be related to the *A. robusta* recovery project, particularly monitoring disturbance and natural regeneration. As can be seen, the experimental design problems allow for rectification to occur, which are a small investment that can guide the recovery efforts’ future direction. Pilot studies can provide a means to refine or better identify the issues associated with species recovery. In the current investigation, refining the experimental design occurred from identifying problems at the pilot stage but more importantly, the identification of new opportunities to aid *A. robusta* recovery occurred.

## Conclusion

Interpreting the results provides some general inferences only but can provide an opportunity to reflect on the experimental design for *A. robusta* recruitment. Comparing *A. robusta* seed germination to a species of least concern would facilitate the species recovery process. The concept is no different from the investigation conducted by Abihudi et al. (2020). Abihudi et al. (2020) demonstrated the benefits of small discrete experimentation on a species could improve species management. *A. robusta* inhabits an environment considered to be prone to disturbance. The pilot investigation identified experimental design shortcomings, which translates to requiring further investigation but focussing on the essential environmental cues. For the recovery of *A. robusta* to be successful, community involvement is essential which can be applied from the lessons learned through experimentation and research at the pilot stage.

